# Microglia-specific promoter activities of human *HexB* gene

**DOI:** 10.1101/2021.10.28.465974

**Authors:** Sahil Shah, Lilly M. Wong, Kendra Ellis, Brittany Bondar, Sami Saribas, Julia Ting, Zhengyu Wei, Yuyang Tang, Xianwei Wang, Hong Wang, Binhua Ling, David M. Margolis, J. Victor Garcia, Wenhui Hu, Guochun Jiang

## Abstract

Adeno-associated virus (AAV)-mediated genetic targeting of microglia remains challenging. Overcoming this hurdle is essential for gene editing in the central nervous system (CNS). Here, we characterized the minimal/native promoter of the *HexB* gene, which is known to be specifically and stably expressed in the microglia during homeostatic and pathological conditions. Dual reporter and serial deletion assays identified the critical role of the natural 5’ untranslated region (−97 bp related to the first ATG) in driving transcriptional activity of the mouse *HexB* gene. The native promoter region of mouse, human and monkey *HexB* located at -135, -134 and -170 bp to the first ATG, respectively. These promoters were highly active and specific in microglia with strong cross-species transcriptional activities, but had no activities in primary astrocytes. In addition, we identified a 135 bp promoter of *CD68* gene was also highly active in microglia but not in astrocytes. Considering that *HexB* is specifically expressed in microglia, not in monocytes/macrophages or other neuronal cells, these data suggest that the newly characterized 134 bp microglia-specific *HexB* promoter can be an ideal candidate for microglia-targeting AAV gene therapy, which could be developed for HIV eradication in the brain wherein microglia harbor the main HIV reservoirs in the CNS.

**Summary:** It is hard to overstate the importance of gene therapy that can remove viral genes from human cells. A cure for HIV would mean a lifetime free of treatment for patients who now must maintain a strict regimen of ART indefinitely. In order to develop a cure using AAV delivery, payload DNA must meet the AAV vector size limitations, and the payload genes must be expressed appropriately. Previous studies have identified microglia-specific *HexB* gene that shows stable expression during neural homeostasis and pathogenesis. Our study identified the essential *HexB* gene promoter (134 bp) as a strong candidate for AAV gene therapy to specifically target the brain microglia, the main cellular reservoirs of HIV in the central nervous system. Our studies continue to move us closer to identifying target-specific gene therapy for NeuroAIDS.

## Introduction

Treatment for HIV-1 (HIV) has become increasingly effective in the past few decades since the epidemic swept the United States. While antiretroviral therapy (ART) decreases viral replication to undetectable levels and improves the quality of life for those affected, there is no cure for HIV/AIDS (1-3). Upon cessation of ART, viral loads will rebound, which often time were even higher than pretreatment levels (3). In the brain, ART drugs have limited bioavailability and may not efficiently cross the blood-brain-barrier (BBB) in the central nervous system (CNS) (4). The establishment of the latent HIV reservoirs in the brain contributes to HIV-Associated Neurocognitive Disorders (HAND), which persists despite ART in people living with HIV (PLWH). In order to eradicate the disease as a whole, cellular HIV reservoirs hosting replication-competent HIV in the brain must be targeted by HIV cure strategies such as gene editing to excise disease-causing genetic material of HIV.

While there is evidence that many cell types harbor HIV DNA and/or RNA in CNS (5, 6), such as brain myeloid cells (BMCs), astrocytes, neural stem cells (7-10), oligodendrocytes and CD4+ T cells (11-15); it remains to be determined which cell type(s) is (are) the major HIV reservoir in CNS. As one major cell type of BMCs, the resident microglia act as the first line of defense against pathogens in CNS (15, 16). They are long-lived (17) and can self-renew *in vivo* (18-23), which may allow the persistence of HIV infection and serve as HIV reservoirs in the CNS. Blood-borne monocyte-derived macrophages (MDMs) are another source of BMCs (about 5-10%) (24, 25). They are in perivascular regions, terminally differentiated and frequently replenished by hematopoietic progenitors. However, with the relative short lifespan (∼ a few months) and lack of self-renewing potential, it is unclear whether CNS MDMs can support long-lasting HIV infection. Mounting evidence supports the notion that the long-lasting HIV reservoirs establish in the CNS, where the brain microglia may constitute the major cellular CNS reservoirs (11, 14, 15, 26-33). In addition, it has been shown that microglia activation and HIV-associated neuronal damage are linked to HAND developed in up to half of PLWH (34-37). Therefore, BMCs, especially microglia, are one of the major targets for HIV curative strategies that would permit ART-free remission and prevent CNS dysfunction.

Discovery of a proper microglia-specific promoter is needed for the development of gene editing tools specific for the eradication of HIV CNS reservoirs. Recent RNAscope and single-cell RNA sequencing (scRNA seq) analysis determined *HexB* is exclusively expressed in mouse brain microglia but not monocytes/macrophages and other neural cells (38, 39). Importantly, this novel microglia-specific gene retains its expression under various pathological conditions, while many other microglia core genes are substantially downregulated (38, 39). This indicates that *HexB* could be an excellent option for microglia-specific targeting gene therapy for CNS injury and diseases. The insert size limitation in most gene therapy vectors has been a main challenge for packaging capacity and transduction efficiency. In this study, we determined whether the *HexB* promoter is specific to microglia and whether its activity is species dependent. We also optimized the essential element of the native human *HexB* promoter that has the potential to be used in the AAV gene therapy for brain HIV eradication.

## Materials and Methods

### Construction of HexB Promoter Vectors

NEBuilder® *HiFi* DNA Assembly cloning kit (NEB, Cat# E5520S) was used to clone *HexB* gene promoters of various sizes and species into a promoterless AAV plasmid vector containing gaussia dura luciferase and GFP (LG) dual reporter. First, PCR from genomic DNA was used to generate *HexB* gene promoter inserts. The three main sources of promoter DNA were human induced pluripotent stem cells (IPS), mouse embryonic stem cells (ES), and monkey LC30 cells. PCR was completed with Phusion High-Fidelity PCR Master Mix kit (Thermo Fisher, F531). At the same time, the promoterless AAV vector was digested with the appropriate restriction enzymes (Thermo Scientific FastDigest) to allow for insertion of the *HexB* promoter. The PCR products were purified from an agarose gel, and the digestion product was purified using Monarch PCR & DNA Cleanup Kit (NEB, T1030S). Following the preparation of both the backbone vector and insert, the two components were ligated with the NEB HiFi assembly system, using a 1:10 ratio of backbone to insert. The assembled vector was then transformed into NEB stable competent cells and incubated overnight. In creating most vectors, colony PCR was used to confirm that the correct promoter size was inserted prior to sequencing. The correct clones were verified by restriction enzyme digestion and Sanger sequencing as well as functional measures.

### Functional Testing of *HexB* gene Promoter in Cell Culture

The first experiments to determine the activity of the *HexB* promoters were performed in HEK293T cells using standard PeiMax transfection method. Then the promoter activities were tested in C20 human microglia and SIM-A9 mouse microglia cell lines using Glial-Mag kit (OZ Bioscience, Cat # GL002500). The cells were cultured under standard conditions and seeded into 96-well plates at a density of 3.0×10^4^/100 μl. HEK293T cells were cultured in DMEM with 10% FBS. C20 cells were cultured in a DMEM with 5% FBS medium. SIM-A9 cells were cultured in DMEM with 10% FBS and 5% horse serum. After growing overnight, the cells were transfected with indicated AAV-HexB vectors. Transfections were completed in quadruplicates where each well received 100 ng of promoter reporter DNA and 20 ng of pSEAP2 control vector for transfection normalization. Empty promoterless LG vector was used as a negative control (empty vector). The CMV-driven pRRL-Flag-LG or pcDNA3-GFP vectors were used as the positive controls. At 24-96 hours post transfection, the cells were assessed for expression *via* fluorescence microscopy of GFP and gaussia luciferase assay.

After preliminary testing in microglial lines, transfections were performed using primary human microglia cells (Celprogen) and primary microglia cells isolated from fresh brain tissue of rhesus macaques. Primary human and NHP microglia were cultured in DMEM/F12 (Gibco) supplemented with 10% defined fetal bovine serum (HyClone), recombinant human M-CSF (R&D Systems), 100 μg/mL streptomycin, 100 U/mL penicillin, 2 mM L-glutamine, 3 mM sodium pyruvate and 10 mM HEPES. Primary human and NHP astrocytes were cultured in astrocyte medium (AM, ScienCell Research Laboratories) supplemented with FBS, P/S and Astrocyte Growth Supplement (AGS) (Sciencell Research Laboratories). All cells were cultured in 37°C incubator with 5% CO2. Cell transfections were carried out in 96-well plate with 1.0×10^4 target cells where each well received 100 ng DNA with a mix of Lipofectamine 3000 (Invitrogen) in Opti-MEM.

### Luciferase Assays and GFP microscopy

Approximately 50 μl of culture supernatants were harvested 24-72 hours post transfection. Prior to luciferase assay, supernatant from cells was combined with an equal volume of coelenterazine (CTZ) substrate that was diluted 1:50 with CTZ dilution buffer. Prior to measurement, samples were allowed to incubate for 5 min at room temperature. Luciferase activity was measured by fluorescence plate reader (5 second integration) per manufacturer’s instructions (NanoLight Technology). SEAP activity in the culture supernatants were measured with QUANTI-Blue™ Solution (Invivogen) following the manufacture’s protocolThe data were analyzed in comparison with control empty vector transfection after SEAP normalization. At the end of transfection, the GFP+ cells were also analyzed by fluorescence microscopy to determine the gene transcription driven by promoters.

### Statistical analysis

Quantification of fold changes in promoter groups compared with corresponding promoterless or promoter groups was performed using excel software. Statistical analysis was performed using Prism GraphPad 9.1. Significance at \**P* < 0.05, ** *P* < 0.01, and *** *P* < 0.001 and **** *P* < 0.0001 was determined using a two-tailed student’s *t*-test between two groups or by one-way ANOVA for multiple comparisons. Data were presented as mean ± SE. The size and type of individual samples were indicated and specified in the figure legends.

## Results

### Validation of mouse *HexB* gene promoter activity with dual reporter assay

Given AAV gene therapy is promising in clinical trials, we selected AAV transfer vector for future application to AAV packaging and gene delivery. We have established dual reporter assay using gaussia dura luciferase (gdLuc) and GFP expression (thereafter referred as LG) that offer several benefits. It can measure the dynamic quantification of secreted luciferase in culture media with highly sensitive gdLuc assay. The transcriptional activity can also be determined by GFP expression with either fluorescence microscopy or flow cytometry (**Fig. 1A**). Both measures are useful for a fast and high-throughput analysis of *HexB* promoter activity. Based on previous reports on the optimal promoter activity of the mouse *HexB* gene (40), we initially evaluated the activity of *HexB* promoter region -330 bp upstream of ATG (mHexB-330) (**Fig. 1A**). We selected HEK293T cells as the test platform because they are highly transfectable. We found that the mHexB-330 robustly facilitated gdLuc reporter expression by 20∼22 fold 48 hours after transfection (**Fig. 1B-C**), which is consistent with a previous report using chloroamphenicol acetyltransferase reporter assay in NIH-3T3 cells (40). Fluorescent microcopy further confirmed the dramatic increase in GFP expression in mHexB-330 transfected cells (**Fig. 1C**). When the 5’-UTR region was deleted, no transcription activity was observed, which validated the critical role of 5’-UTR (−97) in driving the promoter activity of HexB-330 (**Fig. 1B**) (40).

**Figure 1.**
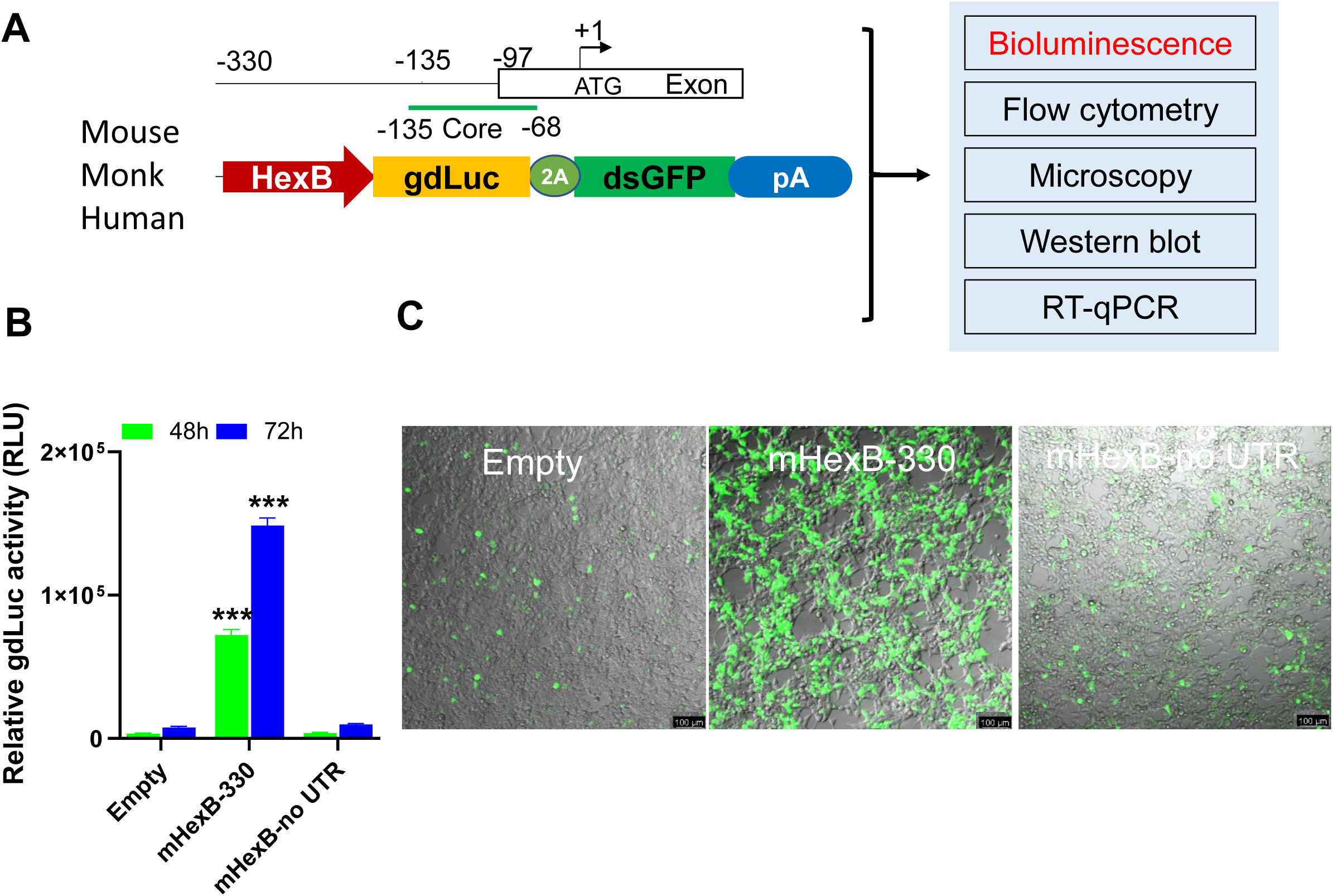
LG dual reporter assays to measure *HexB* promoter activity. (A) Diagram for LG dual reporter assays driven by *HexB* promoter and representative nucleotide numbers for mouse HexB. (B) HEK293T cells were transfected with mouse 330 bp *HexB* promoter-driven dual LG reporter (mHexb-330) and no *HexB* 5’UTR-driven dual LG reporter (mHexB-no UTR) plasmid in 96-well plate, where Empty promoterless LG reporter vector served as negative control. At 48-72 hours after transfection, gdLuc levels in culture media were measured to determine the *HexB* promoter activities. (C) Representative images were taken to show GFP expression in HEK293T cells 72 hours after transfection (scale bar, 100 μm). ***, p<0.001 analyzed with student’s *t*-test compared with empty vector transfection control (n=4).

### Cloning and characterization of essential promoter regions for human and rhesus macaque *HexB*

Because of the high level of sequence conservation across species in the promoter region of *HexB* genes (41), we cloned the essential/minimal promoter region of human *HexB* for the potential application in proviral HIV eradication. As expected, the shorter mouse promoter mHexB-135 (135 bp) retained robust activity, despite moderately weaker than mHexB-330 (**Fig. 2**). Similarly, both human 134 bp *HexB* (hHexB-134) and 183 bp *HexB* (hHexB183) promoters have robust promoter activity, while the shorter promoter (HexB-134) retained high activity in driving gene transcription (**Fig. 2**). However, when a 68 bp core mouse *HexB* promoter (mHexB-Core68) was transfected, no transcription activity was observed compared to an empty vector. This was also true for the human *HexB* promoter, where the 93 bp core (hHexB-Core93) and 43 bp core (HexB-core43) promoters did not have transcriptional activity (**Fig. 2**), indicating the promoter regulatory region (40 bp) at -134 bp to -93 bp is required for human *HexB* promoter activity, consistent with a previous report in the human fibroblast model (42). To expand the potential application to SIV gene therapy in rhesus macaques, we cloned rhesus macaque 170 bp *HexB* (mkHexB-170) promoter (**Fig. 2**), which showed similar activity to the human HexB-183 promoter. These data indicate that the essential and minimal/native *HexB* promoters are sufficient to drive gene transcription.

**Figure 2.**
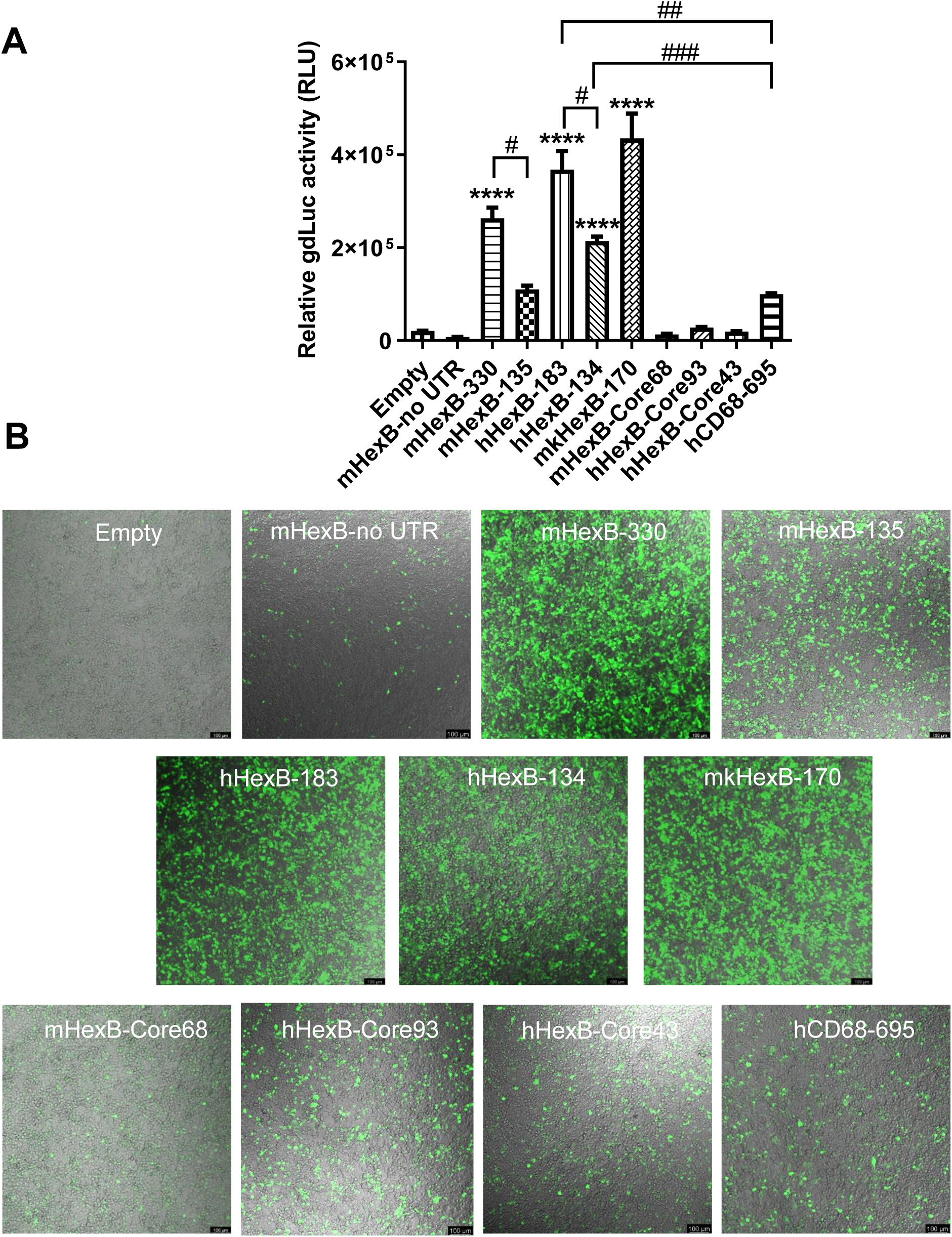
Activity of microglia-related promoters, including *HexB* and *CD68*, in the induction of gene transcription in HEK293T cells. (A) Different sizes of mouse, human or monkey *HexB* promoter-driven dual-LG reporters were transfected into HEK293T cells in 96-well plate. At 72 hours after transfection, gdLuc levels in culture media were measured to determine the *HexB* promoter activity. Cells transfected plasmids driven by other microglia-related promoters, such as *CD68*, were also included, and transcription activity were determined by measuring luciferase levels in culture supernatants. (B) Representative images after transfection are shown (scale bar, 100 μm). ****, p<0.0001, compared with empty vector transfection controls (One-way ANOVA, n=4). #, p<0.05; ##, p<0.01; ###, p<0.001, compared with indicated groups (*t*-test, n=4). m, mouse; h, human; mk, monkey.

### *HexB* promoter is active in microglia cell lines derived from different species

*HexB* is exclusively expressed in microglia in the CNS *in vivo* (39). Thus, we determined whether mouse *HexB* promoter is active in a microglia cell line derived from mouse brain. After transfection, we found that mHexB-330 but not mHexB135 promoter induced luciferase expression in SIM-A9 mouse microglial cells (**Fig. 3A**). hHexB-134, and to a less extent mkHexB-170, was able to drive gene expression in mouse microglia cells. Surprisingly, the longer human HexB promoter, hHexB-183 was less effective to activate transcription, which was similar to the core promoters, hHexB-Core93 and hHexB-Core43.

**Figure 3.**
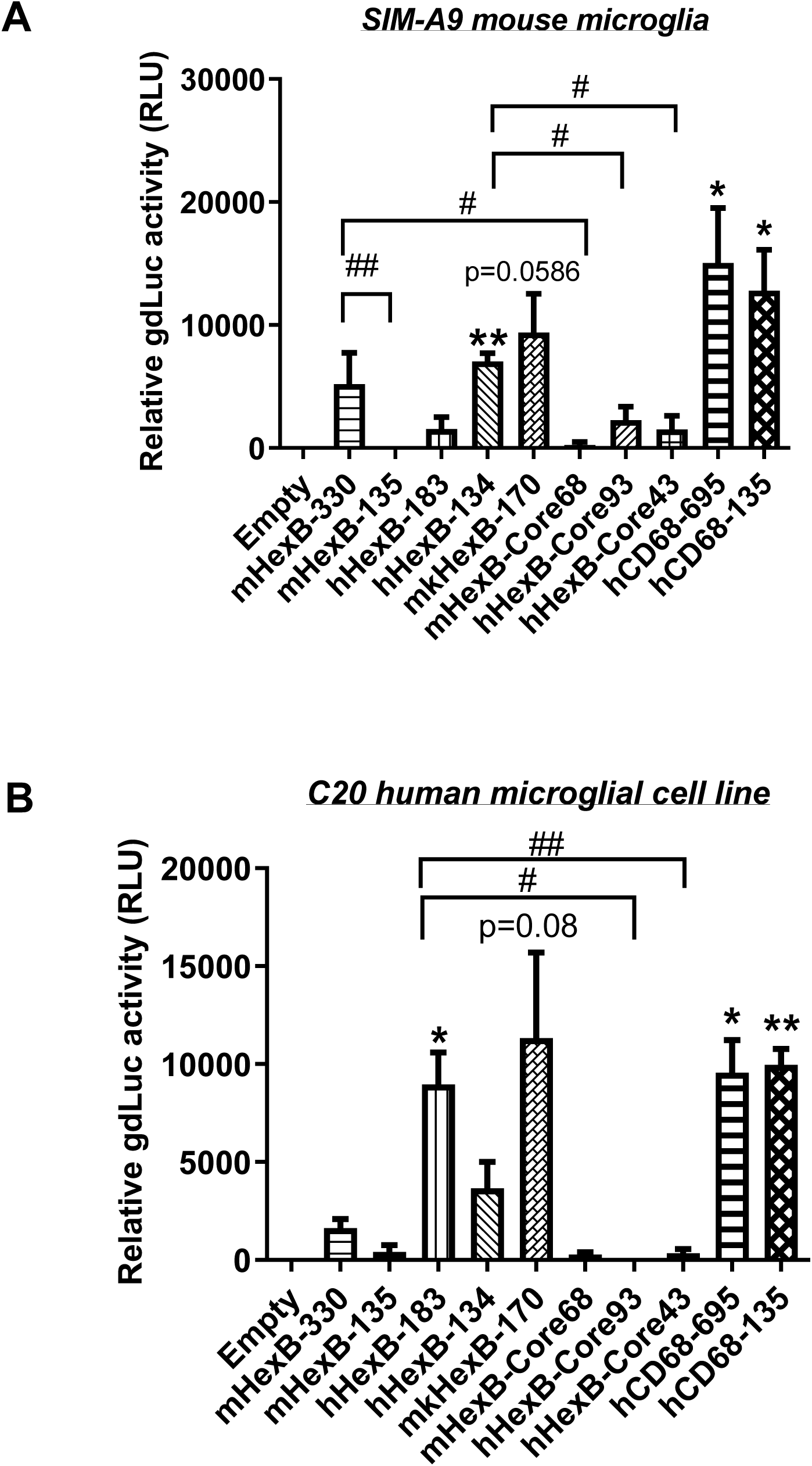
Transcription activity of *HexB* promoters in mouse SIM-A9 and human C20 microglia cells. *HexB* promoter-driven dual LG reporters were transfected into mouse SIM-A9 (A) or human C20 (B) microglial cells. Supernatants were collected 72 hr later and subjected to luciferase assays. *, p<0.05; **, p<0.01; compared with empty vector transfection control (One-way ANOVA, n=4). #, p<0.05; ##, p<0.01, compared with cells with indicated groups (*t*-test, n=4). m, mouse; h, human; mk, monkey.

To see whether there is cross-species activity of *HexB* promoters, we transfected these reporters into human microglia cell line C20. As shown in **Fig. 3B**, mouse HexB promoters (mHexB-330 and mHexB-135) lost their transcriptional capacity. In contrast, both human (hHexB-183) and monkey *HexB* (mkHexB-170) promoters highly induced transcriptional activity, which was lost when the core promoters were transfected into C20 cells. These data showed that human and rhesus macque *HexB* promoters are active in driving gene transcription in both of the microglial cell lines originated from different species, suggesting that *HexB* promoter activity retains a high level of cross-species interaction. It was interesting that the transcription was similarly driven by human or rhesus macaque *HexB* promoters, this may reflect the similar genetic background between human and rhesus macaque. Since mHexB-135 failed to induce transcription in either of the above microglia cell line models (**Fig. 3**), we decided to focus on mHexB-330 promoter in the following studies.

### *HexB* promoter is highly active in driving gene transcription in human and rhesus macaque primary microglia cells, but not in primary astrocytes

While SIM-A9 was derived from cortical tissues collected from postnatal mouse pups and spontaneously immortalized weeks after *in vitro* passaging (ATCC)(43), C20 microglia cell line model was originated from primary human microglia cells transformed by SV40 T antigen and human TERT(44). In order to generate an AAV vector for microglia targeting in a preclinical or clinical setting in the future, it is essential to test its transcriptional activity in the primary human microglia, but not in transformed human microglia. Thus, we examined the promoter activities of these *HexB* promoters in primary microglia cells. We first transfected HexB-driven dual LG reporters into the primary human microglial cells, where a CMV-driven pcDNA3-GFP and CMV-driven dual LG plasmids were included as positive transfection controls. We found that all these *HexB* promoters highly induced GFP expression (**Fig. 4A**). When further quantitated by luciferase assays, we discovered that the transcriptional activities of these *HexB* promoters were strong and comparable with each other (**Fig. 4B**), further supporting the cross-species activity observed in SIM-A9 and C20 (**Fig. 3)**. Surprisingly, when tested in the primary astrocytes, extremely low to no transcription activities were observed in the cells transfected in any species of the *HexB* promoters (**Fig. 4C-D**), either measured by GFP microscopy or by luciferase activity. Also, CMV promoter-driven gene transcription in astrocytes was generally much lower than in microglia. This may be due to that internal transcription factors in astrocytes failed to efficiently support *HexB* promoter activity even though it was transfectable as CMV promoter-driven GFP and luciferase were expressed in astrocytes (**Fig. 4A-D**). It was notable that no luciferase activities were detectable in cells transfected with CMV-driven pcDNA3-GFP plasmids because pcDNA3-GFP does not contain luciferase gene (**Fig. 4A-D**). Similar patterns were observed in primary microglia and astrocytes derived from nonhuman primates, i.e. rhesus macaques (**Fig. 4E-H**), where all these *HexB* promoters actively induces gene expression in microglia cells, but not in astrocytes. Taken together, these data support a notion that *HexB* promoter is able to drive gene transcription in a microglia-specific and possibly cross-species manner.

**Figure 4.**
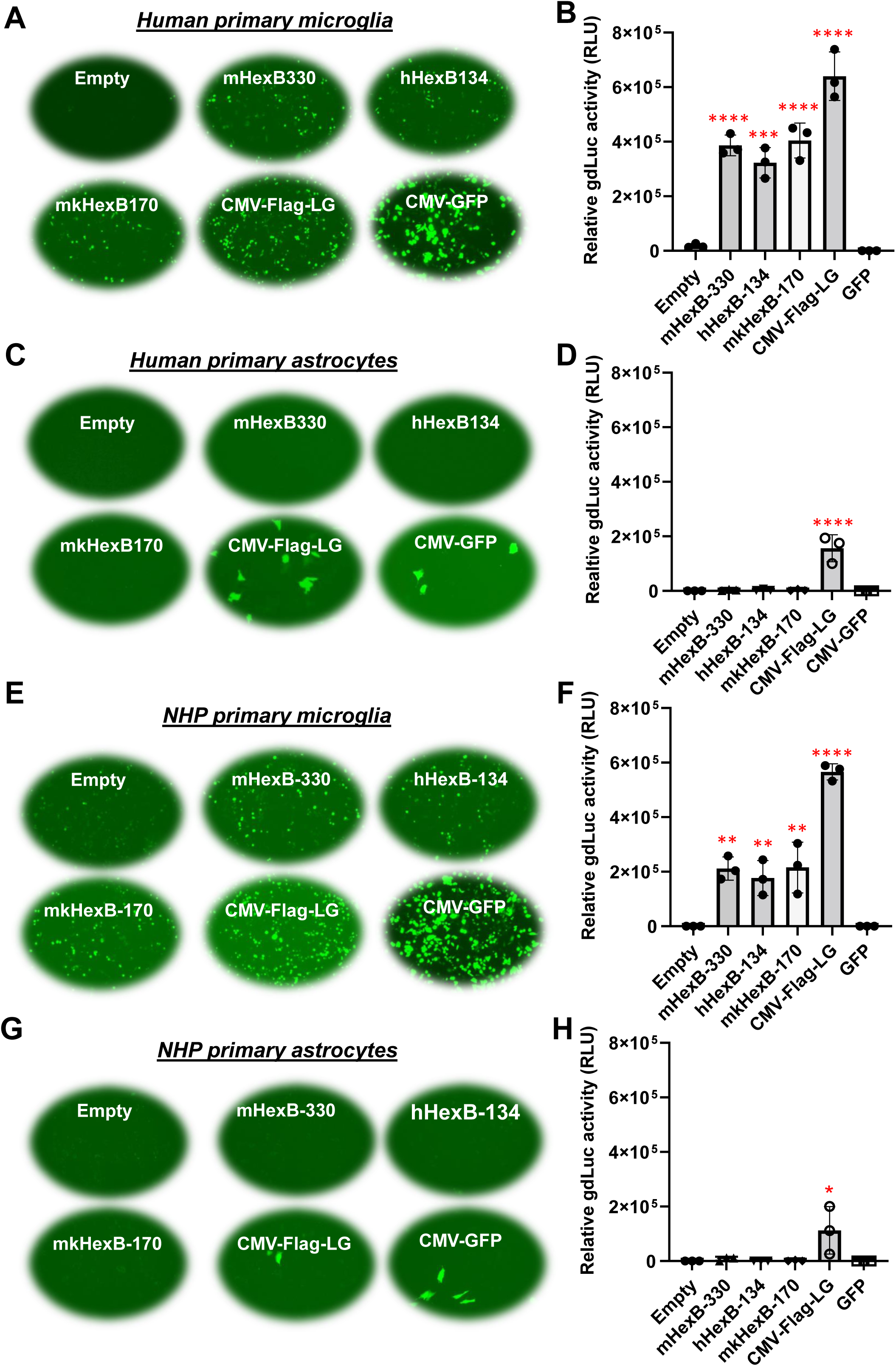
*HexB* promoter induces reporter transcription mainly in microglia and not in astrocytes. Mouse, human or monkey *HexB* promoter-driven dual-LG reporters were transfected into human primary microglia cells (A, B) or human primary astrocytes (C, D) plated in 96-well plates. After transfection, the presence of GFP+ cells was determined by fluorescence microscopy (A and C), and gdLuc levels (B and D) in culture media to determine the activity of the indicated *HexB* promoters. Similarly, *HexB* promoter-driven dual-LG reporters were transfected into primary rhesus macaque microglia (E, F) or primary rhesus macaque astrocytes (G, H) in 96-well plate. After transfection, GFP+ cells were observed with microscopy (E and G, 10x magnification), and gdLuc levels in culture media (F and H) were measured to determine the *HexB* promoter activity. **, p<0.01; ***, p<0.001; ****, p<0.0001, compared with empty vector transfection controls (One-way ANOVA, n=3). m, mouse; h, human; mk, monkey.

### Efficient transcription potency of minimal and essential promoter of human *HexB* gene in microglia cells

*HexB* promoter with a minimal size is important for AAV gene therapy in targeting microglia. In HEK293T cell model, we discovered that the core *HexB* promoters had low to no transcription activities (**Fig. 2**) compared with their native mHexB-330, hHexB-134/183, or mkHexB-170 promoter, regardless of its origin from mouse (mHexB-Core68) or from human (hHexB-Core93/Core43). We have validated these observations in mouse and human microglia cell lines, SIM-A1 and C20 (**Fig. 3A-B**). Like with HEK293T cells, we found that hHexB-core promoters lacked transcription when the functional 5’-UTR were absent (**Fig. 3A-B**). This was similarly observed in the primary microglia cells derived from human or monkey brains (**Fig. 5A-D**). In the primary astrocytes, except CMV-Flag-LG positive control transfection, no transcription was detected in either naÏve or core HexB promoters-transfected cells (**Fig. 5E-H**). These data suggest that, considering the packaging size limits (∼4.8 kb) in AAV packaging, the essential and small-sized human *HexB* promoter hHexB-134 may be an ideal candidate for AAV gene therapy targeting human microglia.

**Figure 5.**
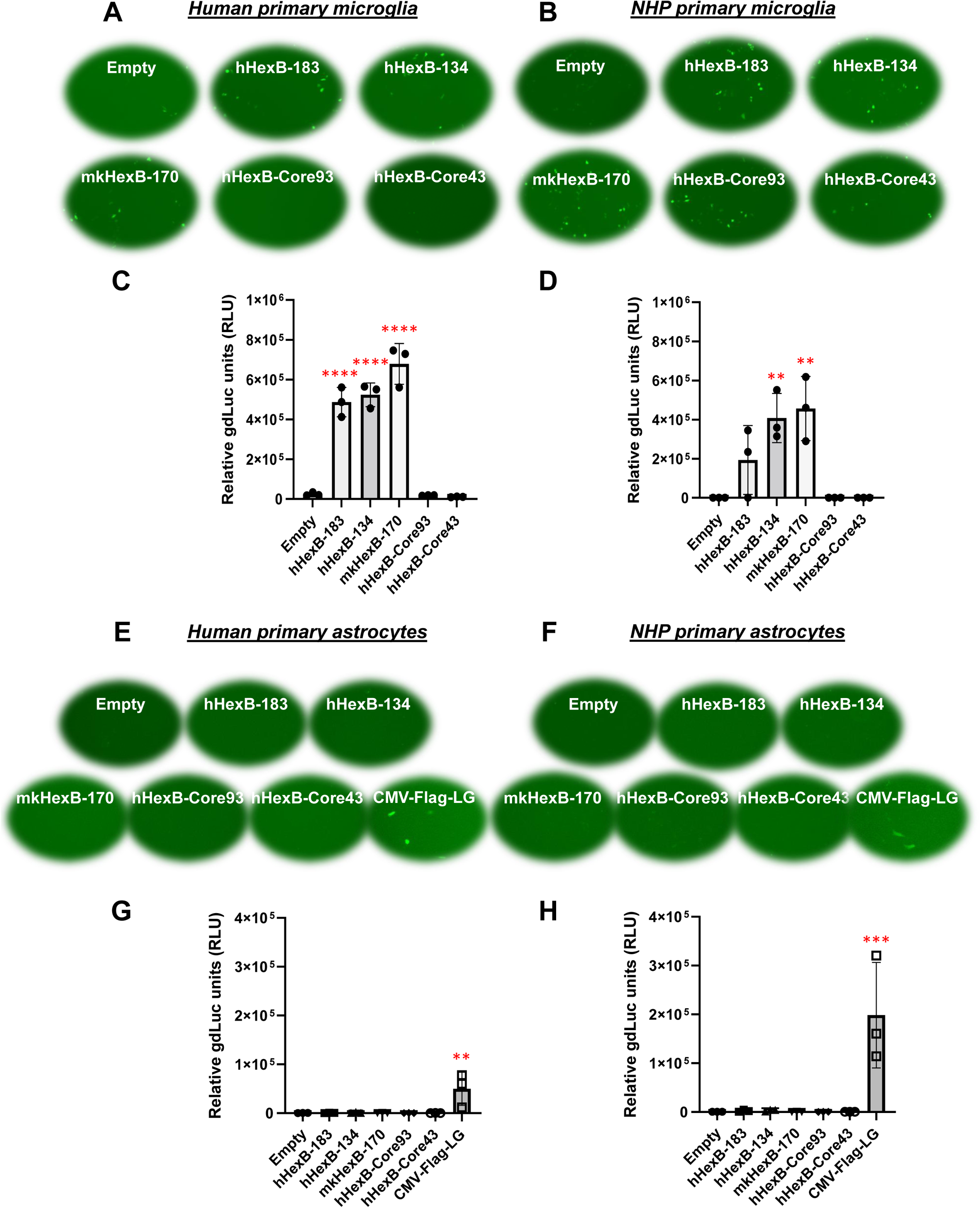
Core *HexB* promoter fails to drive transcription in primary microglia. Dual LG reporter driven by hHexB-183, hHexB-134 or hHexB core promoters were transfected into human (A, C) or monkey (B, D) primary microglial cells in 96-well plate as in figure 4. These plasmids were also transfected into human (E and G) or monkey (F and H) primary astrocytes. At 48 hours post-transfection, GFP+ cells (10x magnification) and gdLuc levels were measured to determine the *HexB* promoter activities. **, p<0.01; ****, p<0.0001, compared with empty vector transfection controls (One-way ANOVA, n=3). m, mouse; h, human; mk, monkey.

### Transcription activity of *HexB* promoter in comparison to other known microglial gene promoters

It has been shown that promoter from myeloid cell markers, such as CD68, were able to drive gene transcription in mouse microglia, which promotes gene targeting in microglia *via* a novel capsid-modified AAV6 vector (45). However, it is not clear whether they have a similar promoter activity in the human microglia, particularly in the primary microglia cells. Since *HexB* promoter displayed unique transcription activity preferred in microglia, it prompted us to compare their capacities of driving gene transcription. We cloned the well-established human *CD68* (695 bp, hCD68-695) promoter and its shorter version of 135 bp promoter (hCD68-135) (16, 38, 44) into our dual LG reporter vector. Initially, we discovered that hCD68-695 showed low transcription activity in HEK293T cells (**Fig. 2**), but strong promoter activities in SIM-A9 and C20 cell lines after transfection (**Fig. 3**). Surprisingly, its short version of counterpart hCD68-135 was also highly active in both microglia cell line models. Since the human *HexB* promoter is more relevant to our human cell study, we set to examine the activities of both promoters in primary microglia. In human primary microglial cells, all these promoters were highly active in gene transcription including hCD68-695 (**Fig. 6A and C**), which was expected. In rhesus macaque primary microglia, hHexB-183 has less promoter activity while hHexB-134, hCD68-135, and hCD68-695 retained high transcription capacity (**Fig. 6B, D**). When tested in the human or rhesus macaque primary astrocytes, no luciferase activity was detectable with any of these microglia-specific promoters (**Fig. 6E-H**). The CMV-Flag LG vector served as positive transfection control (**Fig. 6E-H**). Again, CMV-driven gene transcription in astrocytes was also lower than in microglia (**Fig. 6E-H**). Considering the transcriptional activity and optimal size for AAV packaging, these analyses indicate that hHexB-134 promoter can be further developed to serve as a candidate microglia-specific promoter for AAV gene therapy. Our data also expanded the previous findings of CD68 promoter as an appropriate promoter by identifying a new and much short version of human CD68 (hCD68-135), ideally for gene targeting in microglia cells in the future.

**Figure 6.**
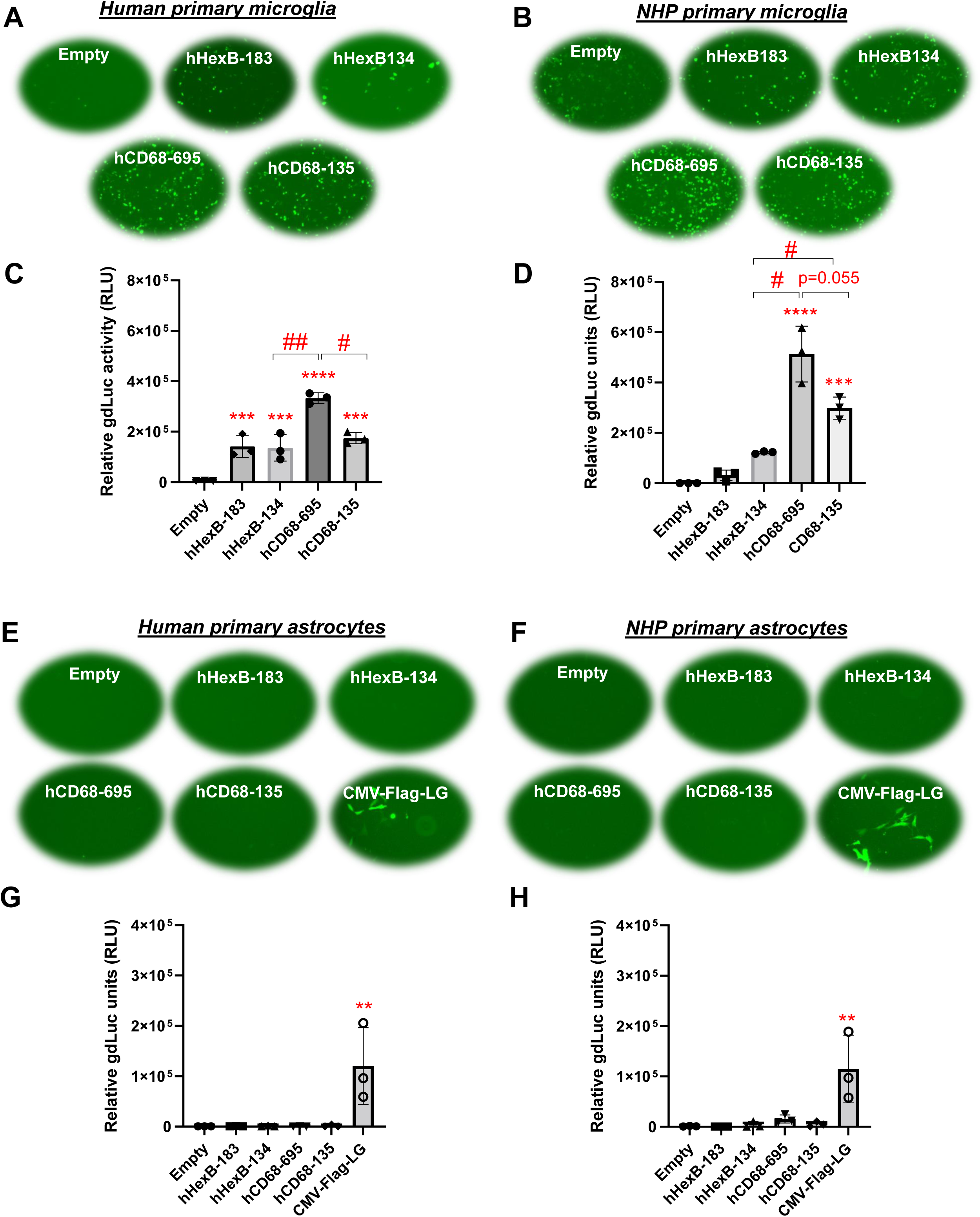
Human *HexB* promoter-induced gene transcription in comparison to other microglia gene promoters in primary human microglia and astrocytes. Microglia-specific promoter-driven dual-LG reporters were transfected into human primary microglia (A and C) or human primary astrocytes (E and G) in 96-well plate. After transfection, GFP+ cells were observed with microscopy (A and E), and gdLuc levels (C and G) in culture media were measured to determine the *HexB* promoter activities. Similarly, promoter-driven dual-LG reporter were transfected into primary rhesus macaque microglia (B and D) or primary rhesus macaque astrocytes (F and H) in 96-well plate. GFP+ cells were observed with microscopy (B and F, 10x magnification), and gdLuc levels in culture media (D and H) were measured to determine the *HexB* promoter activities. *, p<0.05; **, p<0.01; ***, p<0.001; ****, p<0.0001; compared with empty vector transfection controls (One-way ANOVA, n=3). #, p<0.05; ##, p<0.01 (*t-test*, n=3)

## Discussion

As AAV delivery of gene editors as one of the foremost technologies in development for gene therapy of latent HIV reservoirs, finding an effective gene editing system is essential. This is particularly relevant to the HIV cure strategies targeting CNS reservoirs, as the current “shock and kill” strategy may induce high levels of neuron damage or other side effects (46, 47). One way is to develop an effective AAV serotype which can pass through BBB and/or deliver into cellular HIV reservoirs in the brain, such as microglia cells (45). Alternatively, finding an effective promoter to express the selected gene editor specifically in the brain microglia is paramount to progress. Considering the size restrictions for AAV packaging, it is even more important to identify a promoter that is both small and transcriptionally active in microglia. Combination of both approaches may achieve an effective HIV gene editing in the brain *via* not only penetrating BBB but also specific targeting into microglia, one of the major cellular reservoirs in the CNS (13, 48-50).

Microglia-specific gene promoters have been studied in the mouse microglia cell models using mouse myeloid cell-specific promoter including the *CD68* promoter (16, 45); however, it is not clear whether these promoters are effective in human primary microglial cells. CD68 is expressed in myeloid cells, and therefore it is not specific to human microglia. Fortunately, using scRNA-seq technology, recent studies have identified several new biomarkers which have proven specific to brain microglia, including *TMEM119* and *HexB* (38, 39, 44). Among them, *HexB* expression is stable in the microglia *in vivo* even during neuropathogenic conditions, making it a superior marker to for microglia (39, 44).. We found that the *HexB* promoter is specific to microglia cells, possibly in a cross-species manner. Our data support the *HexB* promoter for the development of new gene editing systems targeting microglia (**Figs. 3, 4 and 6**).

Longer endogenous promoters have shown higher levels of gene transcription in the human genome (51). Our studies confirmed that this is crucial in determining if *HexB* promoter is able to induce the GFP and luciferase expression in primary human microglia cells (**Figs. 2 and 5**). In the initial study, mHexB-330, hHexB-183 and mkHexB-170 promoters showed the highest expression of luciferase in a model transformed cell line of HEK293T (**Fig. 2**). When comparing the human *HexB* promoters of sizes 183 bp, 134 bp, 93 bp, and 43 bp, the distinction between the latter two and the former two was clear: the core promoters (93bp and 43bp) were ineffective in boosting luciferase expression over the promoterless vector (Empty) while the longer promoter that includes 5’-UTR was more active to induce transcription. It has been demonstrated that the essential sequences for human *HexB* promoter activity are localized in the region between 150 bp and 90 bp upstream of the ATG codon (40). The pair of promoters evaluated that contain the 5’-UTR (134 bp and 183 bp), had higher transcriptional activity, indicating that this 5’-UTR region is vital for human *HexB* promoter activity. Therefore, the native promoter of *HexB* comprising of the core promoter and the 5’-UTR is essential and minimally required to drive its transcription.

We also investigated the human *CD68* promoters. Initially, we found that 695 bp human *CD68* promoter, hCD68-695, was highly active in SIM-A9 mouse and C20 human microglial cell lines. This was also true in primary human and rhesus macaque microglia (**Figs. 3 and 6**). Surprisingly, the shorter version of the human *CD68* promoter, hCD68-135, was also active in transcription, although less active than the hCD68-695, in microglia (**Figs. 3 and 6**). In comparison to hCD68 promoters, human *HexB* promoters, hHexB-183 and hHexB-134, strongly induced both GFP and luciferase expression in the primary human microglia, which was comparable to the short version of hCD68 promoter hCD68-135, however, slightly lower than the much longer version of hCD68 promoter hCD68-695 (**Figs. 6A and C**).

While our data validated the previous observations of *CD68* promoter activity in mouse microglia, the size of *CD68* promoter is relatively large (695 bp), when compared to the *HexB* promoter (134 bp) or the short version of the hCD68, hCD68-135, which may limit its application in the future. In another report, the *F4/80* (667 bp), *CD68* (543 bp) and *CD11b* (496 bp) promoters were shown to have some activity in AAV-vectors evaluated in microglia cells. However, these promoter are also relatively large (equal or >500 bp) (52) (**Figs. 6A and C**). These observations support that HexB-135 and CD68-135 promoters warrant further investigations for AAV microglia targeting. However, unlike these markers which are mainly expressed in macrophages, HexB does not express in monocytes/macrophages nor in neural cells (38, 39), but exclusively is expressed in brain microglia. This indicates that HexB promoter is advantageous over those promoters for gene therapy in the brain microglia.

Our study also showed that the dual LG reporter assay is powerful in the discovery of novel cell type-specific promoters. Given that this analysis was carried out using a 96-well plate format, it has the potential to be developed into a high throughput assay to screen promoter activity in cell type-dependent and possibly species-dependent manners, which is useful for the gene editing studies in the future.

Taken together, we discovered that the *HexB* gene promoter is a novel tool suitable for microglia specific targeting of gene transfer vectors, which could be exploited for the future development of novel gene editing therapies for the eradication of HIV reservoirs in the CNS.

## ACKNOWLEDGMENTS

This work was supported by Qura Therapeutics, NIMH (R21MH128034) to GJ, and UM1 AI164567 to DMM. BL is supported by R01 MH116844, R01 NS104016, P51OD011104 and P51OD011133, and WH is supported by R01 DA051893 and R01AI145034

## AUTHOR CONTRIBUTIONS

WH and GJ conceived the study. SS, KE, BB, SS, JT, ZW, XW and HW constructed the plasmids and performed the studies in HEK293, SIM-A9 and C20 microglia models. LMW and YT performed studies involved in human and rhesus macaque primary microglia and astrocytes. BL, DMM and JVG provided necessary advice during the discussion of the study. WH and GJ wrote the manuscript with multiple revisions. All authors read and approved the manuscript for submission.

